# Apo- and holo- transferrin differentially interact with ferroportin and hephaestin to regulate iron release at the blood-brain barrier

**DOI:** 10.1101/2023.01.10.522344

**Authors:** Stephanie L. Baringer, Kondaiah Palsa, Ian A. Simpson, James R. Connor

## Abstract

**Background:** Apo- (iron free) and holo- (iron bound) transferrin (Tf) participate in precise regulation of brain iron uptake at endothelial cells of the blood-brain barrier. Apo-Tf indicates an iron deficient environment and stimulates iron release, while holo-Tf indicates an iron sufficient environment and suppresses additional iron release. Free iron is exported through ferroportin, with hephaestin as an aid to the process. Until now, the molecular mechanism of apo- and holo-Tf’s influence on iron release was largely unknown.

**Methods:** Here we use a variety of cell culture techniques, including co-immunoprecipitation and proximity ligation assay, in iPSC-derived endothelial cells and HEK 293 cells to investigate the mechanism of apo- and holo-Tf’s influence over iron release. We placed our findings in physiological context by further deciphering how hepcidin played a role in this mechanism as well.

**Results:** We demonstrate that holo-Tf induces the internalization of ferroportin through the established ferroportin degradation pathway. Furthermore, holo-Tf directly binds to ferroportin, whereas apo-Tf directly binds to hephaestin. Only pathological levels of hepcidin disrupt the interaction between holo-Tf and ferroportin, and no amount of hepcidin disrupts the interaction between apo-Tf and hephaestin. The disruption of the holo-Tf and ferroportin interaction by hepcidin is due to hepcidin’s ability to rapidly internalize ferroportin compared to holo-Tf.

**Conclusions:** These novel findings provide a molecular mechanism for apo- and holo-Tf regulation of iron release from endothelial cells. They further demonstrate how hepcidin impacts these protein-protein interactions, and offer a model for how holo-Tf and hepcidin corporate to suppress iron release. We have established a more thorough understanding of the mechanisms behind iron release regulation with great clinical impact for a variety of neurological conditions in which iron release is dysregulated.

## Background

Regulation of iron uptake at the blood-brain barrier (BBB) is crucial for proper brain function. Detrimental alterations in brain iron homeostasis can lead to a variety of neurological conditions, including but not limited to neurodegenerative diseases (Alzheimer’s disease, Parkinson’s disease, and amyotrophic lateral sclerosis)^1^ and Restless Legs Syndrome^2^. Our group and others have shown that endothelial cells (ECs) of the BBB serve as reservoirs for iron before it is subsequently released into the extracellular fluid of the brain. Moreover this release is regulated by levels apo (iron free)- and holo (iron bound)-transferrin (Tf)^3–7^ in extracellular fluid. Using both *in vitro*^3,4,6^ and *in vivo*^7^ models, we have shown that increasing the ratio of apo-to holo-Tf, reflecting an iron deficient environment, stimulates iron release from ECs, whereas elevated holo-Tf relative to apo-Tf, reflecting an iron-replete environment, suppresses iron release. This feedback mechanism allows for regional specificity of iron uptake based on regional iron consumption and metabolic needs^8,9^.

Free iron is released from cells, including ECs, through ferroportin (Fpn), the only know iron exporter. Fpn function is aided by a number of proteins, including hephaestin (Heph)^10,11^, a ferroxidase that converts ferrous (Fe2+) to ferric (Fe3+) iron. Heph is required for both the stability of Fpn in the plasma membrane and the efflux of iron through Fpn^10,11^. Inversely, Fpn can be inhibited by hepcidin^12^, a pro-inflammatory peptide hormone, primarily secreted by the liver^13^ and in small amounts by astrocytes^14^. When hepcidin binds to Fpn, Fpn is ubiquitinated for internalization and subsequent degradation^12,15^. Simpson *et al*. found that, in addition to iron release, holo-Tf also decreases Fpn protein in EC culture models of the BBB^6^ but the mechanism is unclear. Conversely, it has been proposed that apo-Tf participates in interactions with ferroxidases such as Heph and ceruloplasmin to facilitate iron release^16–18^. In the present study, we have determined the differential interactions that apo- and holo-Tf have with Fpn and Heph to control iron release. Moreover, we demonstrate the impact that hepcidin can have on these interactions. By understanding the regulatory mechanism of iron release into the brain, numerous neurological diseases with iron uptake dysregulation can be better studied and potentially treated.

## Methods

### Cell Culture

Human endothelial-like cells (ECs) were differentiated from ATCC-DYS0100 human iPSCs as described previously^19,20^. Briefly, iPSCs were seeded onto a Matrigel-coated plate in E8 medium (Thermo Fisher Scientific, 05990) containing 10µM ROCK inhibitor (Y-27632, R&D Systems, 1254) at a density of 15,000 cells/cm^2^. The iPSCs differentiation was initiated by changing the E8 medium to E6 medium (Thermo Fisher Scientific, A1516401) after 24 hrs seeding. E6 medium was changed daily up to 4 days before switching to human endothelial serum free medium (hESFM) (Thermo Fisher Scientific, 11111) supplemented with 10nM bFGF (Fibroblast growth factor, Peprotech, 100-18B) and 10 µM all-trans retinoic acid (RA, Sigma, R2625) and 1% B27 (Thermo Fisher Scientific, 17504-044). After 48 hrs of no medium changes, cells were harvested and replated onto Transwell filters coated with collagen IV and fibronectin. Twenty-four hours after replating, bFGF and RA were removed from the medium to induce barrier phenotype. HEK 283 cells were maintained in Dulbecco’s modified Eagle’s medium (DMEM, Gibco, 11965-084) and supplemented with 10% FBS and 1% penicillin-streptomycin (Gibco, 15070063).

### Proximity Ligation Assay (PLA)

PLA is a technique that precisely demonstrates if two proteins directly interact with one another. When two proteins are in close enough proximity to be interacting, the secondary oligomer probes ligate together, allowing for the amplification of the oligomers and resulting in a fluorescent signal. PLA was performed using a Duolink assay kit (Sigma-Aldrich, DUO92013) according to the manufacturer’s instructions^21^. Chamber slides (Falcon, 354108) were coated with poly-D-lysine 2 hrs before HEK 293 cells were culture on the slides at a density of 15,000 cell/cm^2^. In order to remove an exogenous Tf, 24 hrs later the media was replaced with DMEM containing no FBS. Cells were exposed to apo-or holo-Tf (Sigma, T1147 and T4132) for 10 minutes and then washed to procced with PLA. PLA was performed the following day. Primary antibodies used were the following: myelin basic protein 1 (MBP1, Abcam, ab22460, 1:500), ferritin (Abcam, ab77127, 1:500), Tf (ProteinTech, 66161-1, 1:500), TfR (Cell Signaling, 13208S, 1:500), Tf (Abcam, ab82411, 1:500), Fpn (gift from M. Knutson, 1:500), and Heph (Santa Cruz, SC-365365, 1:500). Positive and negative controls used for assay optimization can be found in Supplemental Figure 2. Imaging and analysis were performed using Revolve R4 microscope (Echo). The integrated density was calculated by summing the pixels from PLA signal and dividing by the field of view area. The integrated density of background from negative controls were subtracted from these values. To determine the integrated density per cell, this was then divided by the number of cells in the field of view. A minimum of three images were taken in different regions of the slides and then averaged for a single biological replicate. Image brightness was uniformly increased for the purposes of publication but not for quantification.

### Plasmid and Transfection

HEK 293 cells were seeded at a density of 7 × 10^4^ cell/cm^2^ in a 6-well plate. The following day, the cells were transfected with 1μg/well of the HA-tagged Fpn plasmid (Vector Builder, VB220407-1185gaa, Supplemental Figure 1) using Lipofectamine™ 3000 Transfection Reagent (Invitrogen, L3000001).

### Co-immunoprecipitation

In order to remove an exogenous Tf, the media was replaced with DMEM containing no FBS 24 hrs before the start of experiments. Cells were exposed to apo-or holo-Tf (Sigma, T1147 and T4132) for 10 minutes and then washed on ice with cold PBS twice. Chilled 100μl Co-IP lysis buffer (20 mM Tris HCl, pH 8, 137 mM NaCl, 10% glycerol, 1% Triton x-100, and 2 mM EDTA) was added to each well. Cells were collected and incubated with rotation for 30 minutes at 4°C. Cell solutions were centrifuged at 14,000 x g for 20 minutes at 4°C. Supernatant was collected, and protein estimation was performed using Pierce BCA Protein Assay Kit (Thermo, 23227). Approximately 1 mg of protein was used for Co-IP using anti-HA magnetic beads (Thermo, 88837) or Protein G magnetic beads (Thermo, 10003D) complexed with anti-Heph antibody (Santa Cruz, SC-365365) according to manufacturer’s instructions^22^. Briefly, magnetic beads were washed twice with PBS before adding lysates. The bead and lysate solutions were incubated with rotation for 30 minutes at room temperature. After washing with PBS, protein was eluted from beads by resuspending in non-reducing sample buffer and boiling at 90°C for 10 minutes. Magnet was used to isolate the magnetic beads from the protein solution, which was then reduced using 2 M DTT and then loaded for immunoblotting.

### Membrane Protein Isolation

Cells were washed with PBS three time before incubating with 200μl digitonin buffer (20mM Tris-HCl, 250 mM sucrose, 0.007% digitonin, 1x protease inhibitor cocktail)^23^. Cells were gently lifted from the plate and collected in chilled glass mini homogenizers. Once homogenized, samples were spun at 1,500 x g for 10 minutes. The pellet was reserved and the supernatant was spun again at 10,000 x g for 10 minutes. The resulting pellet was combined with the pervious pellet and resuspended in RIPA buffer and 1x protease inhibitor cocktail. After immunoblotting was performed on the samples, the membranes were stained for total protein content using Ponceau S staining solution (Thermo, A40000279) to use as a loading control.

### Immunoblotting

Samples were loaded onto a 4-20% Criterion TGX Precast Protein Gel (Bio-Rad)^7^. Protein was transferred onto a nitrocellulose membrane and probed for Fpn (Alpha Diagnostics, MTP11-S, 1:1000), DMT1 (Millipore, ABS983, 1:1000), Heph (Santa Cruz, SC-365365, 1:1000), TfR (Santa Cruz, sc-65882, 1:250), Tf (Abcam, ab82411, 1:1000), HA tag (Invitrogen, MA5-27915, 1:1000), or cyclophilin B (Abcam, ab16045, 1:1000) as a loading control. Corresponding secondary antibody conjugated to HRP was used (1:5000, GE Amersham) and bands were visualized using ECL reagents (Perkin-Elmer) on an Amersham Imager 600 (GE Amersham). Cellular lysate samples were normalized to cyclophilin B protein as a loading control, and then subsequently normalized to an untreated control sample within each experiment. Membrane protein samples were stained with Ponceau S and normalized to total protein as a loading control.

### Statistical Analysis

Statistical analyses were performed using Prism 9.2 software (Graphpad Software Inc.). Data from at least three independent biological replicates were averaged and are expressed as the mean ± standard error of the mean (SEM). One-way ANOVA with Tukey post-hoc analysis, two-way ANOVA with Sidak’s post hoc analysis, or unpaired t-tests were used to evaluate for statistical significance where appropriate. A p-value <0.05 was considered significant.

## Results

### Holo-Tf decreases Fpn levels through Fpn’s degradation pathway

In the first series of experiments, we examined the effects of apo- and holo-Tf on the cellular levels of Fpn by incubating iPSC-derived ECs with increasing concentrations of either apo-or holo-Tf in hESFM for 8 hours. ECs were cultured onto Transwell inserts and apo-or holo-Tf was placed in the basal chamber to represent the brain-side. The ECs were collected and probed for various iron transport proteins. Incubations with holo-Tf decreased Fpn protein levels by 50% at concentrations as low as 0.1 μM (*p<0.05, Fig. 1A) whereas apo-Tf had no impact on Fpn (Fig. 1A). Other iron transport proteins, such as Heph, DMT1, and TfR, were unchanged with incubations of apo-or holo-Tf (Supplemental Figure 3).

**Figure 1:**
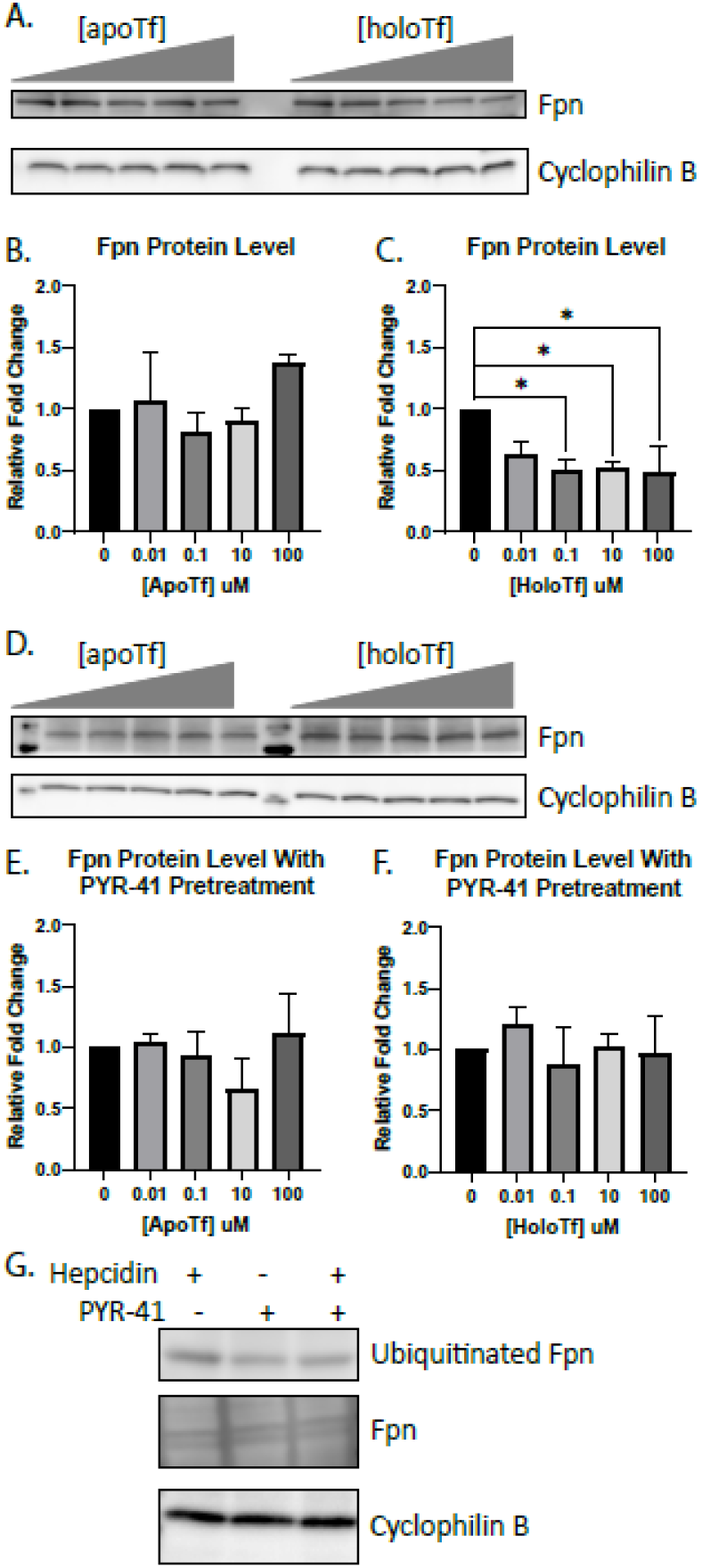
Modulation of Fpn protein levels in ECs by holo-Tf. iPSC-derived ECs were cultured on bi-chamber plates, incubated with apo- or holo-Tf in the basal chamber, and collected after 8 hours for immunoblotting. Fpn protein levels were normalized to cyclophilin B as a loading control. All quantifications were further normalized to untreated control to account for cell count variability. Holo-Tf decreased Fpn protein levels by 50% at concentrations as low as 0.1 μM, while apo-Tf did not (**A-C**). Holo-Tf-mediated internalization and degradation of Fpn was inhibited by a ubiquitination inhibitor, PYR-41, (**D-F**) confirming that holo-Tf’s decreases Fpn through the established degradation pathway. PYR-41’s inhibition of ubiquitination was validated using hepcidin to induce Fpn ubiquitination (**C**). Exposure to hepcidin alone for 30 minutes increases ubiquitination of Fpn. When pretreated with PYR-41 for an additional 30 minutes, this increase of ubiquitination of Fpn is blocked. Total Fpn levels are unchanged. n=3 for all experiments, means of biological replicates ± SEM were evaluated for statistical significance using one- way ANOVA with Tukey’s posttest for significance. *p<0.05

The degradation pathway for Fpn involves ubiquitination by E1 ubiquitin ligase, resulting in the internalization and degradation of Fpn^15,24^. To determine if this classic degradation pathway was the cause of the decreased Fpn induced by holo-Tf, we pretreated ECs with 50 μM PYR-41, an E1 ubiquitin ligase inhibitor, before exposure to either apo-or holo-Tf. The inhibition of Fpn ubiquitination resulted in a mitigation of holo-Tf’s decrease of Fpn (Fig. 1D), while apo-Tf continued to have no impact on Fpn levels (Fig. 1D). Hepcidin, a known inducer of Fpn ubiquitination, was used as a positive control to confirm the function of PYR-41’s inhibition (Fig. 1G). ECs were exposed to 500nm of hepcidin following pretreatment with 50 μM PYR-41 for 30 minutes. Controls were either solely exposed to hepcidin or PYR-41. Hepcidin alone increased Fpn ubiquitination and PYR-41 pretreatment prevented this increase (Fig. 1G).

### Apo- and holo-Tf differentially interact with Fpn and Heph

We next aimed to determine if holo-Tf interacted directly with Fpn. We used HEK 293 cells transfected with an HA-tagged Fpn plasmid to selectively pull-down HA-Fpn. We incubated the cells with 0.25 μM of either apo-or holo-Tf (physiological level in CSF^25^) in media containing no FBS for 10 minutes prior to co-immunoprecipitation (co-IP). Regardless if the cells were incubated with either apo-or holo-Tf, Tf was co-immunoprecipitated with HA-Fpn (Fig. 2A). This indicates that apo- and holo-Tf bind to the Fpn complex of proteins. Because Heph aids Fpn in the export of iron^11^, we hypothesized that apo-Tf could bind to Heph, leading to its co-immunoprecipitation with HA-Fpn. To confirm this, we incubated ECs, which have greater Heph expression than HEK 293 cells, with either apo-of holo-Tf, and performed co-IP with Heph antibody Again, in cells incubated with either apo-of holo-Tf, Tf was co-immunoprecipitated (Fig. 2B) further confirming that Fpn, Heph, and Tf complex together.

**Figure 2:**
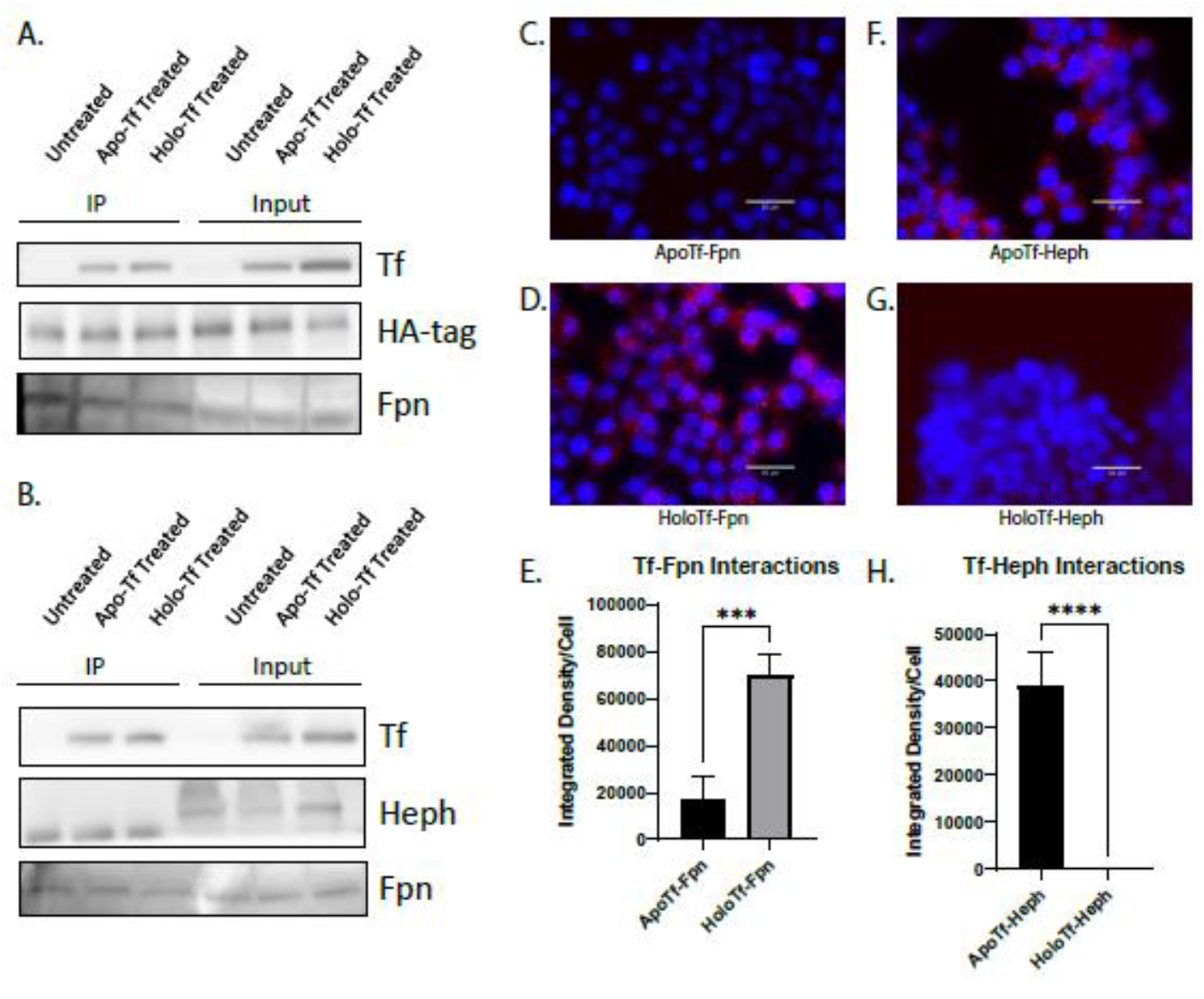
Apo- and holo-Tf interactions with Fpn and Heph. HEK 293 cells were transfected with HA-tagged Fpn and subsequently incubated with 0.25 μM apo- or holo-Tf. Immunoprecipitate (IP) and 50% of cell lysate (input) was processed for immunoblotting. Co-IP of HA-Fpn shows that both apo- and holo-Tf are pulled down along with the Fpn complex (**A**). Co-IP of Heph in iPSC-derived ECs replicated these data (**B**). HEK 293 cells were used to determine direct protein interactions using PLA, reported as integrated density per cell in the field of view per image. Holo-Tf interacts with Fpn (**D**), while apo-Tf does not (**C**). Alternatively, apo-Tf interacts with Heph (**F**), while holo-Tf does not (**G**). n=4 for all experiments, means of biological replicates ± SEM were evaluated for statistical significance using unpaired t test. ***p<0.001, ****p<0.0001

Because co-IP precipitates the entire complex of Fpn, Heph, apo-Tf, and holo-Tf, we aimed to better differentiate if apo- and holo-Tf directly interact with Fpn and Heph by employing proximity ligation assay (PLA), a highly sensitive method of detecting protein-protein interactions. HEK 293 cells were incubated with 0.25 μM of either apo-or holo-Tf in media containing no FBS for 10 minutes. Cells incubated with holo-Tf showed PLA signal when probing for a Tf and Fpn interaction (Fig. 2D). While cells incubated with apo-Tf showed PLA signal when probing for a Tf and Heph interactions (Fig. 2F). Thus, holo-Tf directly interacts with Fpn while apo-Tf does not (***p<0.001, Fig. 2E). Conversely, apo-Tf directly interacts with Heph, while holo-Tf does not (****p<0.0001, Fig. 2H).

### High levels of hepcidin interrupt the interaction between holo-Tf and Fpn

Hepcidin is a well-known regulator and binding partner of Fpn, therefore we aimed to understand how the novel interaction between holo-Tf and Fpn could be impacted by physiological conditions that contribute to iron release. To do so, we used PLA to examine if hepcidin competed with holo-Tf for binding to Fpn. HEK 293 cells were co-incubated with 500 nM hepcidin and varying concentrations of holo-Tf (Fig. 3A-F). Hepcidin interrupted the interaction between 0.25 μM holo-Tf and Fpn (Fig. 3D), resulting in an 75% reduction of PLA signal (*p<0.05, Fig. 3F) compared to no hepcidin treatment (Fig. 3A). Hepcidin was able to reduce the PLA signal by nearly 90% when the concentration of holo-Tf was only 0.025 μM (**p<0.01, Fig. 3E, F). When holo-Tf was present in higher concentrations (25 μM and 2.5 μM), hepcidin did not interrupt the interactions between holo-Tf and Fpn (Fig. 3B, C) but these concentrations of holo-Tf are likely supraphysiological.

**Figure 3:**
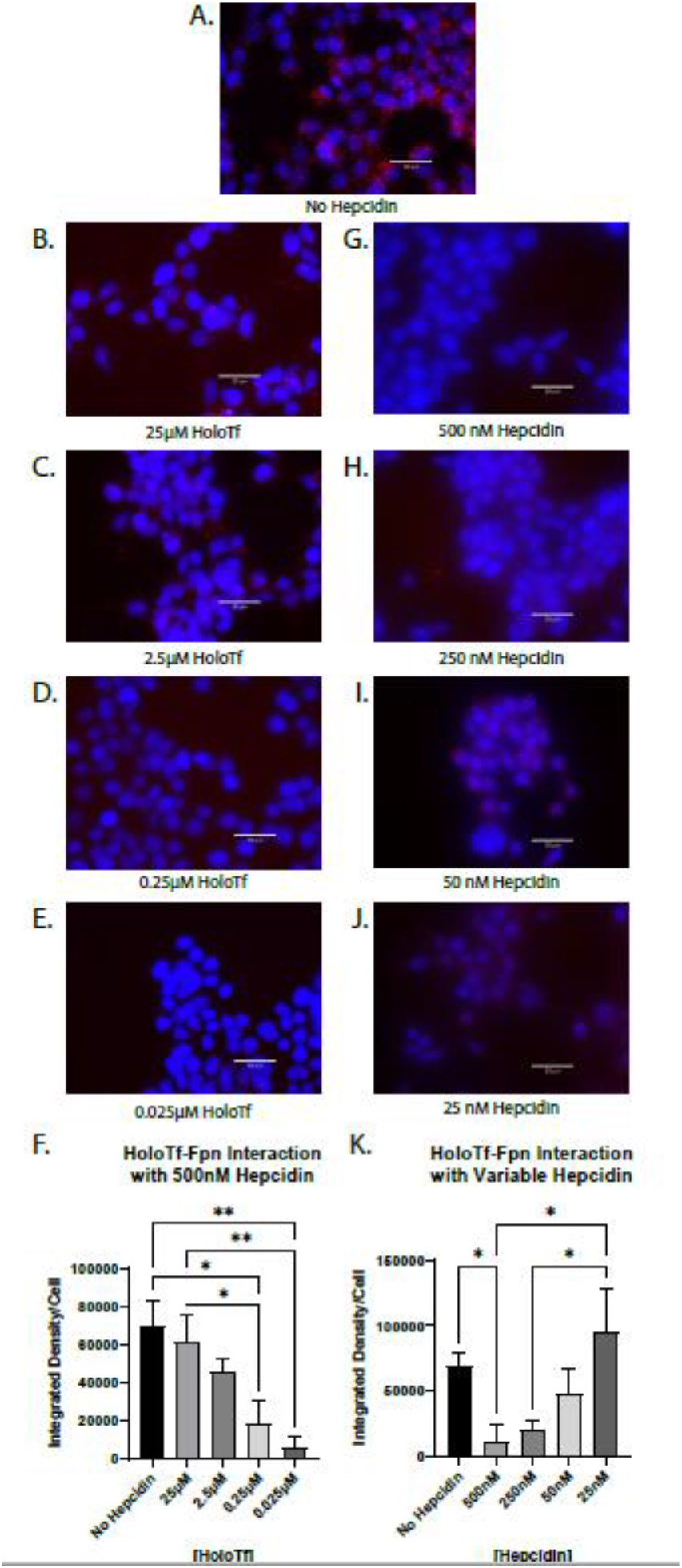
Hepcidin impact on interaction between holo-Tf and Fpn. HEK 293 cells were used to determine the impact of hepcidin on holo-Tf and Fpn interactions using PLA, reported as integrated density per cell in the field of view per image. High levels of hepcidin interrupt the interaction between holo-Tf and Fpn when holo-Tf is present in physiological levels (**D and G**), but not when holo-Tf concentrations are higher (**B and C**) or hepcidin concentrations are closer to baseline physiological (**H-J**). n=3 for all experiments, means of biological replicates ± SEM were evaluated for statistical significance using one-way ANOVA with Tukey’s post-test for significance. *p<0.05, **p<0.01

To determine if the amount of hepcidin was crucial to the interruption of the holo-Tf and Fpn interaction, we performed the reverse competition experiment and co-incubated HEK 293 cells with 0.25 μM holo-Tf and varying concentrations of hepcidin (Fig. 3G-K). Hepcidin interrupted the interaction between holo-Tf and Fpn in a dose dependent manner. Only the highest concentration of 500 nM significantly interrupted the interaction between holo-Tf and Fpn (*p<0.05, Fig. 3G, K). As the concentration of hepcidin decreased, the PLA signals for holo-Tf and Fpn interactions, increased (Fig. 3G-J). The physiological baseline concentration of hepcidin^26^, 25 nM, had no impact on the holo-Tf-Fpn interaction (Fig. 3J).

### Hepcidin does not interrupt the interaction between apo-Tf and Heph

Apo-Tf has been shown to stimulate iron release despite the presence of hepcidin^4^, thus we hypothesized that hepcidin would have no impact on the interaction between apo-Tf and Heph using PLA. HEK 293 cells were co-incubated with 500 nM hepcidin and varying concentrations of apo-Tf (Fig. 4B-F). Unlike with holo-Tf, 500 nM hepcidin did not interrupt the interaction between any amount of apo-Tf and Heph (Fig. 4B-E), as indicated by the unchanged PLA signal (Fig. 4F). In the reverse competition experiment, we co-incubated HEK 293 cells with 0.25 μM apo-Tf and varying concentrations of hepcidin and indicated in (Fig. 4G-J). Again, no concentration of hepcidin was sufficient to alter the interaction between apo-Tf and Heph (Fig. 4K).

**Figure 4:**
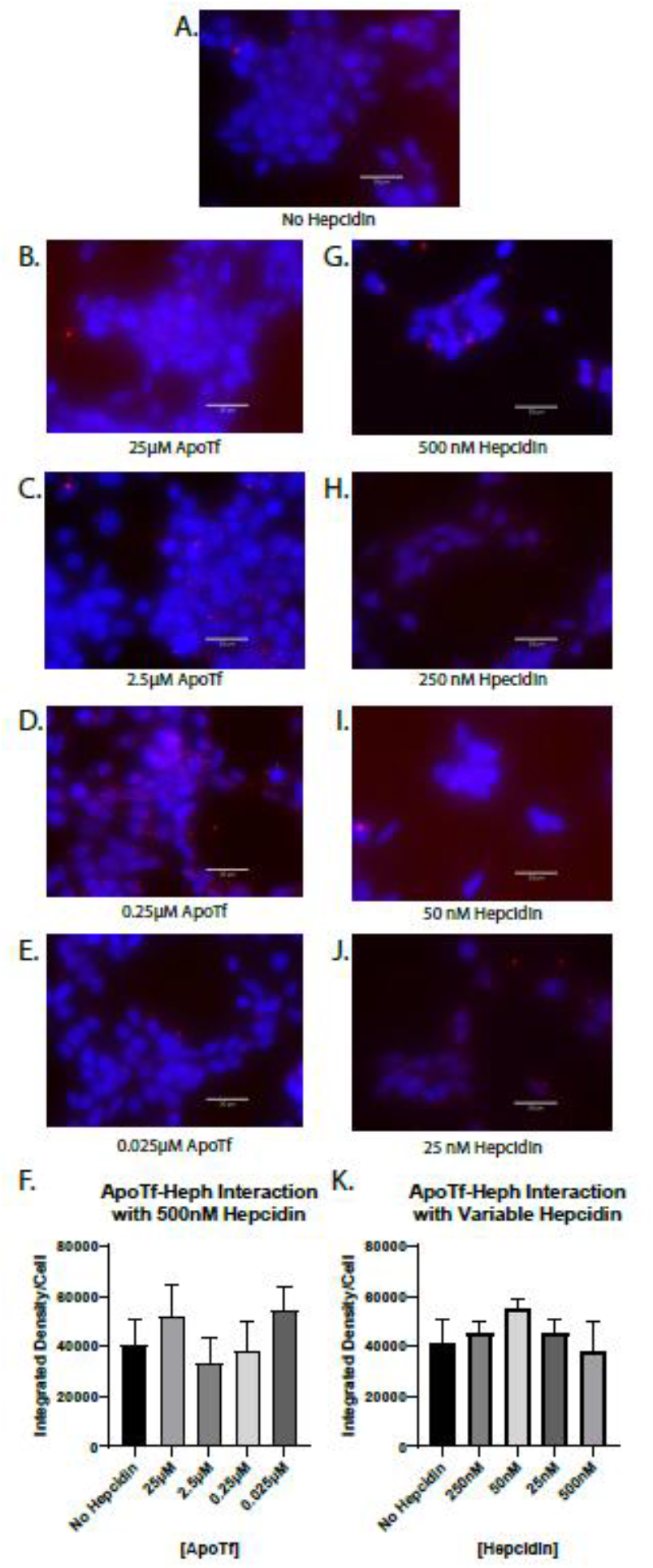
Hepcidin impact on interaction between apo-Tf and Heph. HEK 293 cells were used to determine the impact of hepcidin on apo-Tf and Heph interactions using PLA, reported as integrated density per cell in the field of view per image. Hepcidin has no impact on the interaction between apo-Tf and Heph at any apo-Tf concentrations (**B-E**) or at any hepcidin concentrations (**G-J**). n=3 for all experiments, means of biological replicates ± SEM were evaluated for statistical significance using one-way ANOVA with Tukey’s post-test for significance.

### Hepcidin internalizes Fpn faster than holo-Tf

The PLA experiments showed there was competition between holo-Tf and hepcidin, but did not differentiate if this was due to hepcidin directly competing with holo-Tf for a binding site on Fpn or by internalizing Fpn faster than holo-Tf. To answer these questions, we utilized pretreatment with PYR-41, which prevents the degradation of Fpn and thus removes internalization dynamics as a factor in the binding of holo-Tf and hepcidin to Fpn. We performed PLA on HEK 293 cells exposed to 0.25 μM holo-Tf alone (Fig. 5A), 0.25 μM holo-Tf and 500 nM hepcidin (Fig. 5B), and pretreatment of 50 μM PYR-41 and then 0.25 μM holo-Tf and 500 nM hepcidin (Fig. 5C). As before, hepcidin interrupts the interaction between holo-Tf and Fpn (*p<0.05), however, this decrease in interaction is prevented when with the PYR-41 pretreatment (***p<0.001, Fig. 5A-D). This finding indicates that hepcidin decreases the interaction between holo-Tf and Fpn due to its ability to rapidly internalize Fpn. We further confirmed a decrease of Fpn membrane presence by isolating membrane bound proteins for immunoblotting (Fig. 5E-F). The co-incubation of 0.25 μM holo-Tf and 500 nM hepcidin results in a significant decrease of membrane Fpn protein (*p<0.05, Fig. 5E-F). This decrease in membrane Fpn is prevented when pretreated with PYR-41 (*p<0.01, Fig. 5E-F). These data align with the PLA results and suggests that hepcidin prevents holo-Tf from binding to Fpn by inducing the rapid internalization of Fpn.

**Figure 5:**
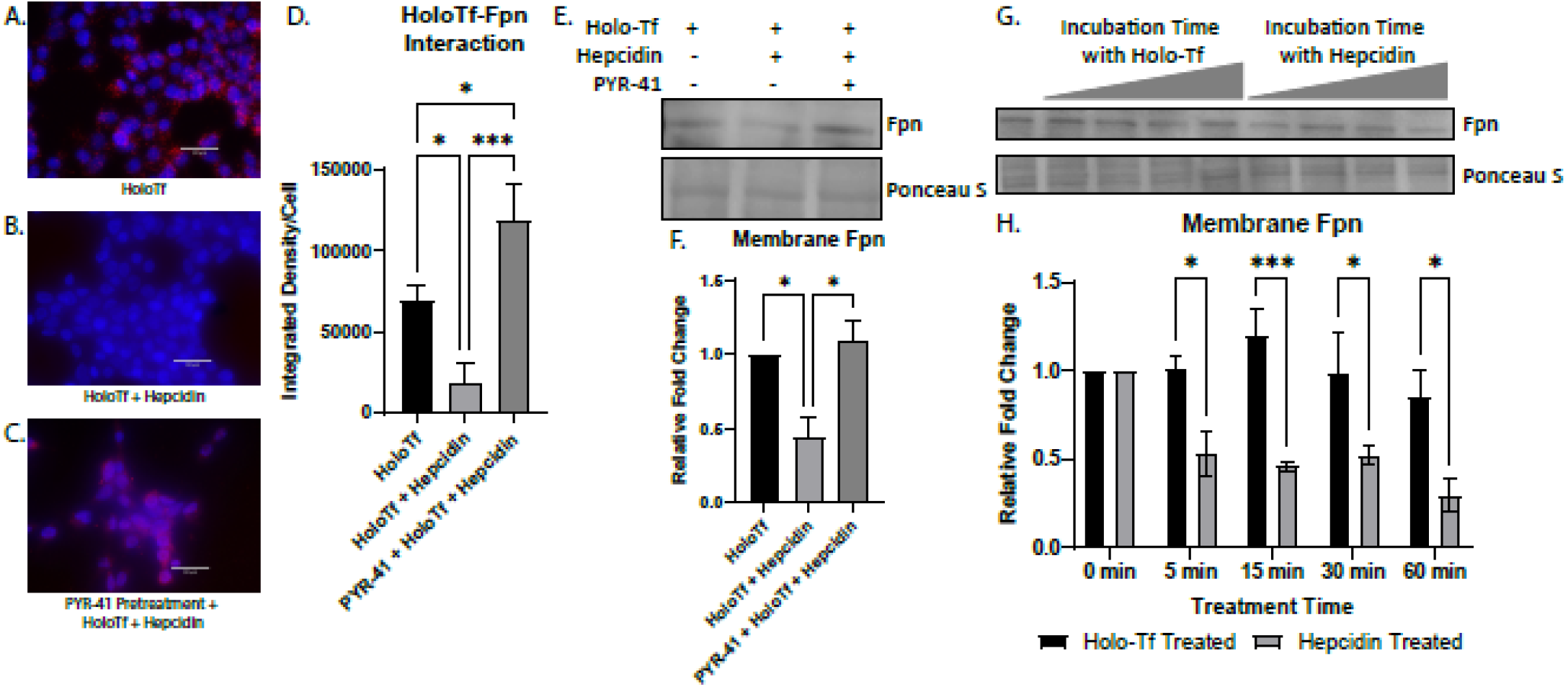
Modulation of Fpn internalization by hepcidin and holo-Tf. HEK 293 cells were used to determine the dynamics of holo-Tf and hepcidin on Fpn internalization using PLA, reported as integrated density per cell in the field of view per image (**A-D**). Pretreatment with PYR-41 (**C**) prevented the hepcidin induced reduction of interaction between holo-Tf and Fpn (**B**). The isolation of membrane bound Fpn confirms that hepcidin and holo-Tf co-incubation greatly reduces membrane Fpn levels, and this is prevented with PYR-41 (**E-F**). Hepcidin reduces membrane Fpn at a faster rate than holo-Tf (**G-H**). n=3 to 5 for all experiments, means of biological replicates ± SEM were evaluated for statistical significance using one-way ANOVA with Tukey’s posttest for significance (**D**) and (**F**) or two-way ANOVA with Sidak’s post-test for significance (**H**). *p<0.05, ***p<0.001

The rate of Fpn internalization induced by holo-Tf or hepcidin was further examined by incubating HEK 293 cells with either 0.25 μM holo-Tf or 500 nM hepcidin over time and subsequently isolating the membrane bound proteins. After only 5 minutes of 500 nM hepcidin incubation, membrane Fpn levels were decreased by nearly 50% compared to holo-Tf treatment (*p<0.05, Fig. 5G-H). The trend continues at incubation times of 15 minutes (***p<0.001), 30 minutes (*p<0.05), and 60 minutes (*p<0.05, Fig. 5G-H). By 60 minutes, hepcidin has internalized 70% of membrane Fpn compared to holo-Tf (*p<0.05, Fig. 5G-H). It is only at 60 minutes that holo-Tf starts to internalize Fpn, with about a 20% decrease compared to 0 minutes (Fig. 5G-H).

## Discussion

This study addresses the molecular mechanisms by which apo- and holo-Tf regulate iron release at the BBB. More specifically, this study demonstrates that apo- and holo-Tf differentially interact with Heph and Fpn. Through its interaction, holo-Tf reduces Fpn protein levels, and this is through Fpn’s established degradation pathway as shown when Fpn degradation is inhibited. Holo-Tf directly interacts with Fpn as shown by orthogonal techniques. Furthermore, when incubated together, hepcidin can interrupt this interaction at high levels that correspond with inflammation or high systemic iron levels, but not at levels that correspond with baseline levels. Hepcidin’s interruption is likely due to its ability to internalize Fpn faster than holo-Tf and not due to direct competition for the same binding site, as we additionally demonstrate herein. On the other hand, hepcidin does not interrupt the interaction between apo-Tf and Heph. These findings offer a glimpse at the mechanism of free iron release into the brain, a crucial process for neurological health.

Fpn is the only known iron exporter, thus control of membrane Fpn is control of free iron release. The internalization and subsequent degradation of Fpn has been extensively studied in the context of hepcidin^12,15,24,27^. Briefly, once hepcidin binds to Fpn, it triggers the ubiquitination of the Fpn, thus signaling for its internalization and lysosomal degradation. Simpson *et al*. showed that by incubating BRECs with 12.5 μM holo-Tf, the levels of Fpn decreased^6^. Here we have replicated those findings in iPSC-derived ECs but at a physiological level; transferrin is found in CSF at about 2 mg/dL, or 0.25 μM^25^. We demonstrate that a basal incubation of as low at 0.1 μM holo-Tf results in a 50% decrease of membrane Fpn. These data provide a mechanistic explanation for why holo-Tf suppresses iron release from ECs. What’s more, other iron-related proteins, such as Heph, DMT1, and TfR, are unchanged with basal holo-Tf exposure. Interestingly, even when exposed to high amounts of holo-Tf, the levels of Fpn do not decrease beyond 50%, suggesting there is a plateaued effect of holo-Tf within the 8-hour experimental time window. The holo-Tf-mediated internalization of Fpn is blocked when the ubiquitination of Fpn is inhibited, suggesting that holo-Tf exerts its effect through the established degradation pathway, similar to hepcidin.

To complete the process of iron export, Fpn works in a complex with many proteins, including Heph^10,11^. Heph is a ferroxidase that converts the Fpn-exported ferrous (Fe2+) to ferric (Fe3+) that can bind to apo-Tf and be utilized by cells. Numerous studies have shown that Heph is required to stabilize Fpn in the plasma membrane and to enable iron export^10,11,28,29^. We have replicated these findings, by demonstrating that Fpn and Heph can be co-immunoprecipitated from ECs. Furthermore, we demonstrate the novel finding that both apo- and holo-Tf independently are co-immunoprecipitated with Fpn and Heph. These results suggest that apo- and holo-Tf bind to Fpn and Heph in a complex of iron export proteins. In order to narrow down which protein holo-Tf bound to in the membrane that resulted in decreasing Fpn, we employed PLA. We found that holo-Tf directly interacts with Fpn, while apo-Tf does not. On the other hand, apo-Tf interacts with Heph, while holo-Tf does not, a finding that is supported in the literature^16,18,30^. It is hypothesized that apo-Tf binds to Heph to accept the ferric iron that Heph converts from ferrous iron. This stimulates the release of more iron as long as there is apo-Tf to accept it. Taken together these data suggest that apo- and holo-Tf differentially interact with iron export proteins, likely due to their structural differences^31^. The exact binding sites, conformation changes, and catalysts for these interactions are an exciting unexplored area that could pave the way for clinical manipulation. For example, as has been done experimental^7^, Tf could be infused to modulate iron accumulation in diseases in which it is dysregulated. Additionally, pharmaceuticals could be designed to facilitate or inhibit the endogenous protein interactions in an effort to correct brain iron accumulation.

Prior to the discovery that elevated holo-Tf could suppress iron release, hepcidin was the primary focus of iron release regulation^13^. Hepcidin is a pro inflammatory hormone peptide primarily secreted by the liver and upregulated in environments of inflammation and high iron levels^32^. Astrocytes^33,34^ and the choroid plexus^35,36^ have also been shown to secrete hepcidin, though in much smaller amounts that cannot account for total brain hepcidin levels^36,37^, suggesting much of the brain hepcidin comes from systemic levels when pathologically necessary, though this has not yet been proven. A number of groups have shown that astrocytic hepcidin reduces Fpn levels and subsequent iron release^14,38,39^. However, we have previously demonstrated that supraphysiological levels of hepcidin are not capable of blocking iron release from ECs^3,4^. These data suggest that hepcidin cannot be the sole regulator of iron release in the brain. In support of this notion, Enculescu *et al*. modeled iron levels, and when compared to their experimental results, the study found that hepcidin control over iron uptake was necessary, but not sufficient^40^. Once a secondary regulatory mechanism was added to the model, their experimental results aligned with the model^40^. Thus, our data directly support that hepcidin is not the sole regulator of iron release and indicate the additional regulators are apo- and holo-Tf.

Our data offer an opportunity to explore the concept of regulation of iron uptake in general by hepcidin. We found that hepcidin competes with holo-Tf for binding to Fpn at low holo-Tf and high hepcidin concentrations. However, when there was more holo-Tf or less hepcidin present, this effect was reduced. Notably, when hepcidin was only present at physiological baseline levels^26^, there was no interruption of the interaction between holo-Tf and Fpn. These findings suggest that hepcidin is only effective at controlling Fpn levels at levels consistent with inflammation or high iron. In observing competition between holo-Tf and hepcidin for Fpn binding, the internalization of Fpn was inhibited to determine if the competition was for binding site availability or rate of internalization. By preventing the internalization of Fpn, hepcidin had no impact on the interaction between holo-Tf and Fpn. This suggests that hepcidin internalizes Fpn faster than holo-Tf, which was confirmed by isolating membrane Fpn. Hepcidin reduces membrane Fpn by nearly 50% in 5 minutes, whereas holo-Tf only starts to reduce membrane Fpn at 60 minutes. On the other hand, no amount of hepcidin impacts the interaction between apo-Tf and Heph. These data offer the intriguing suggestion that if apo-Tf is present, it will bind to Heph even in pathological states and may be an explanation for iron accumulation in neurodegenerative disease. It has been postulated that in Alzheimer’s disease^41^ and Parkinson’s disease^42^ the brain may start as functional iron deficient, along with elevated levels of apo-Tf, which triggers increased iron uptake until the excess iron detrimentally damages the BBB and surrounding cells. The question remains however, if the binding of apo-Tf to Heph will continue to stimulate iron release in the presence of hepcidin.

The model of apo- and holo-Tf regulation of iron release from ECs works as a feedback loop. As cells, such as neurons or astrocytes, need iron for metabolic processes, myelin synthesis, or dopamine synthesis, they take up holo-Tf through TfR^43^. Once endocytosed, the iron is removed and the resulting apo-Tf is released^43^. The communication of brain iron status via apo- and holo-Tf allows cells to signal their iron needs based on their iron consumption. Numerous studies have shown higher regional iron uptake that correspond to areas with higher iron needs^9,44,45^. Our pervious data suggest that as the apo- to holo-Tf ratio changes in the extracellular fluid, more iron is released locally from the BBB. In support of this notion are data showing CSF from iron deficient monkeys and iron chelated astrocytes increase iron release from cultured bovine retinal ECs (BRECs), while iron loaded biological samples resulted in decreased iron release^6^. These data have been replicated when cells are exposed to apo- or holo-Tf directly^3,4,6^ or when apo- or holo-Tf is directly infused into the brain^7^. In all studies mentioned here, apo-Tf increased iron release while holo-Tf decreased iron release.

The data in this study expand the model for brain iron uptake by suggesting that apo-Tf stimulates iron release by binding to Heph to access exported free iron (Fig. 6A). Once loaded with iron, the now holo-Tf becomes available to surrounding cells. If the levels of holo-Tf in the extracellular fluid rise, holo-Tf binds to Fpn to suppress more iron release (Fig. 6B). The internalization of Fpn by holo-Tf is not rapid, unlike hepcidin. When upregulated and present in high amounts, hepcidin can rapidly internalize Fpn (Fig. 6C). Thus, we propose that hepcidin is likely used as a fast acting, immediate stop to iron release in environments of inflammation and very high iron. However, for moment-by-moment regional control of iron release, holo-Tf may be a better candidate to regulate regional iron supply

**Figure 6:**
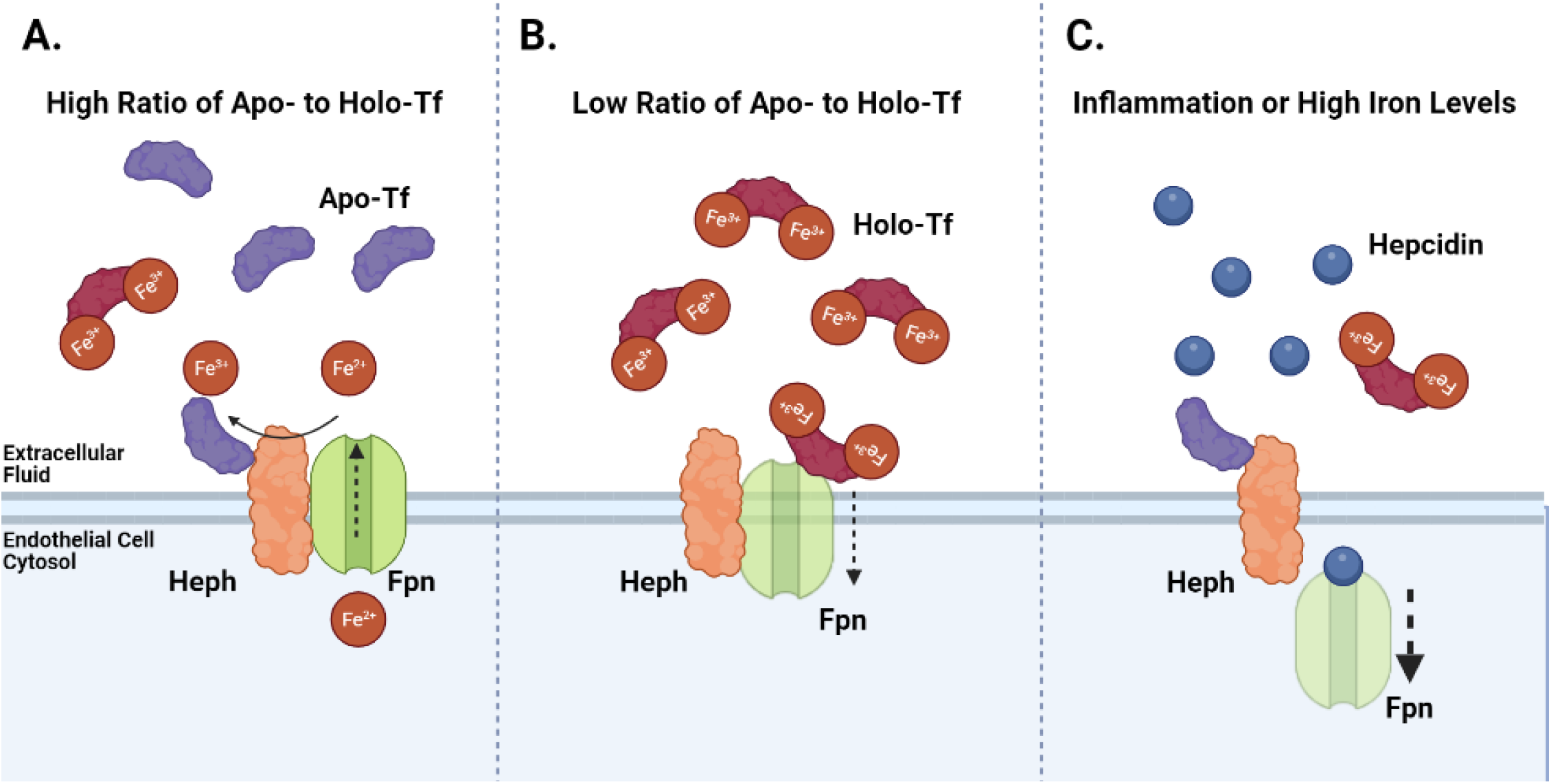
Model of Iron Release Regulation. In our proposed model, in areas that have higher ratios of apo- to holo-Tf (**A**), apo-Tf binds to Heph in order to accept the exported free iron and further stimulates iron release through Fpn. Alternatively, areas of lower ratios of apo- to holo-Tf (**B**), excessive holo-Tf binds to Fpn to facilitate the internalization and degradation of Fpn, and thus suppressing iron release through Fpn. In environments of inflammation or high iron levels, hepcidin production is upregulated (**C**). Hepcidin binds to Fpn and rapidly triggers Fpn’s internalization and abruptly stops free iron release.

## Conclusions

The regulation of brain iron uptake is not influenced by systemic levels^46^, thus a local source is needed. The data herein provide insights into a local regulatory process. This study is the first demonstration that apo- and holo-Tf differentially interact with Fpn and Heph to regulate iron release from ECs of the BBB. Moreover, we have identified a physiologically relevant dynamic between hepcidin and holo-Tf and their influence on membrane Fpn levels. Hepcidin interrupts the interaction between holo-Tf and Fpn by internalizing Fpn much faster than holo-Tf. Furthermore, we show that hepcidin does not interrupt the interaction between apo-Tf and hepcidin. These data suggest the mechanism of free iron release from ECs at the BBB and provide opportunity for further studies in neurological disease models to understand how this mechanism may be disrupted in each disease.

## Abbreviations

(iron free transferrin): Apo-Tf
(BBB): Blood-brain barrier
(co-IP): Co-immunoprecipitation
(ECs): Endothelial cells
(Fpn): Ferroportin
(Heph): Hephaestin
(iron bound transferrin): Holo-Tf
(iPSCs): Induced pluripotent stem cells
(PLA): Proximity ligation assay
(Tf): Transferrin
(TfR): Transferrin receptor

## Declarations

### Ethics approval and consent to participate

Not applicable.

## Consent for publication

Not applicable.

## Availability of data and materials

Data sharing is not applicable to this article as no datasets were generated or analyzed during the current study.

## Author Contributions

SLB and KP performed experiments using iPSC-ECs. SLB performed remaining experiments. All authors contributed to designing research, analyzing data, and writing the paper. All authors read and approved the final manuscript.

## Acknowledgments

The authors would like to thank Drs. Spiegelman and Elcheva and Stem Cell and Regenerative Program for their support, Dr. Ethan Lippmann for providing protocols and guidance to culture and differentiate iPSC-derived EC-like cells, and Dr. Mitch Knutson for providing the ferroportin antibody used for PLA assays. Research reported in this publication was supported by the National Center for Advancing Translational Sciences of the National Institutes of Health Award Number TL1TR002016. The content is solely the responsibility of the authors and does not necessarily represent the official views of the NIH.

## Supplemental Data

**Supplemental Figure 1.**
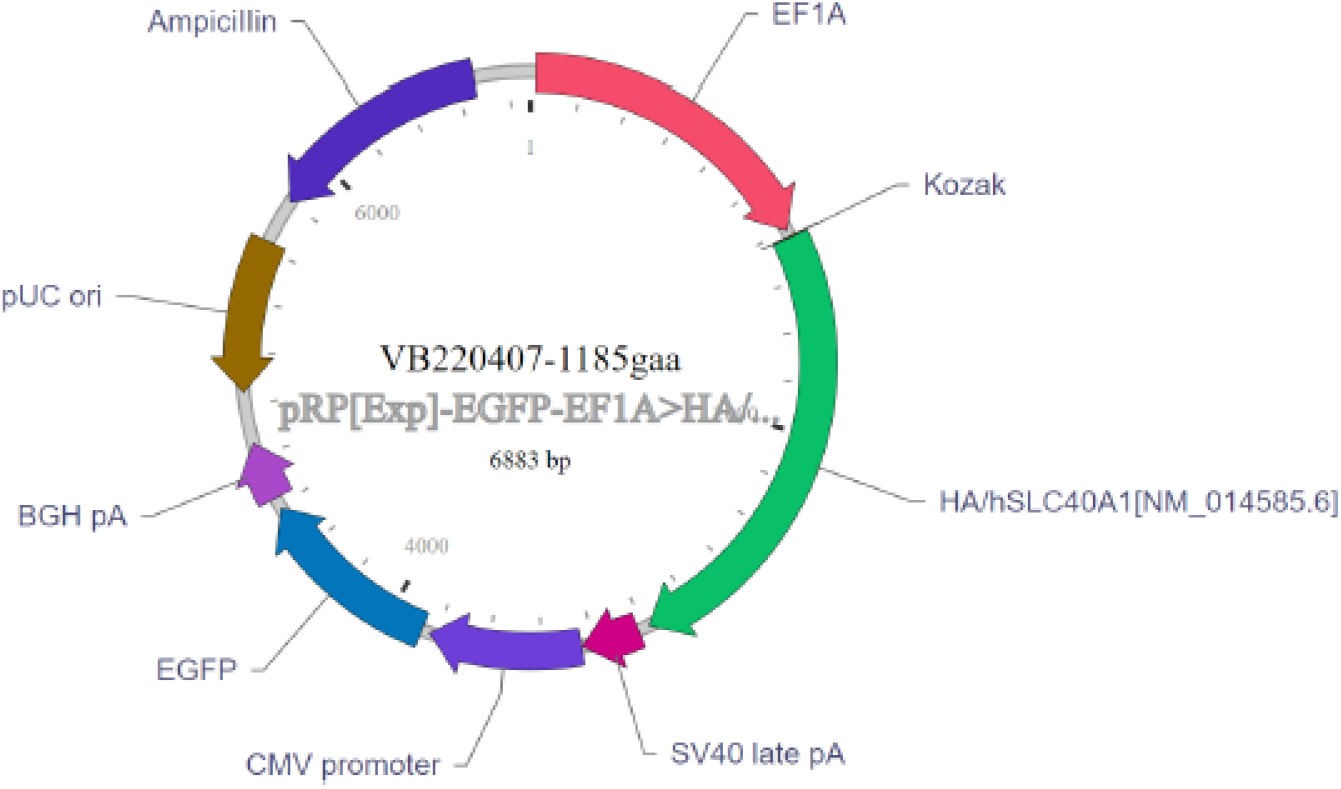
HA-tagged Fpn Plasmid Map. HEK 293 cells were transfected with an HA-tagged Fpn plasmid in order to effectively pull down Fpn in co-IP experiments. The plasmid was designed using Vector Builder. The full sequence, as well as purchasing options, are available online https://en.vectorbuilder.com/vector/VB220407-1185gaa.html.

**Supplemental Figure 2.**
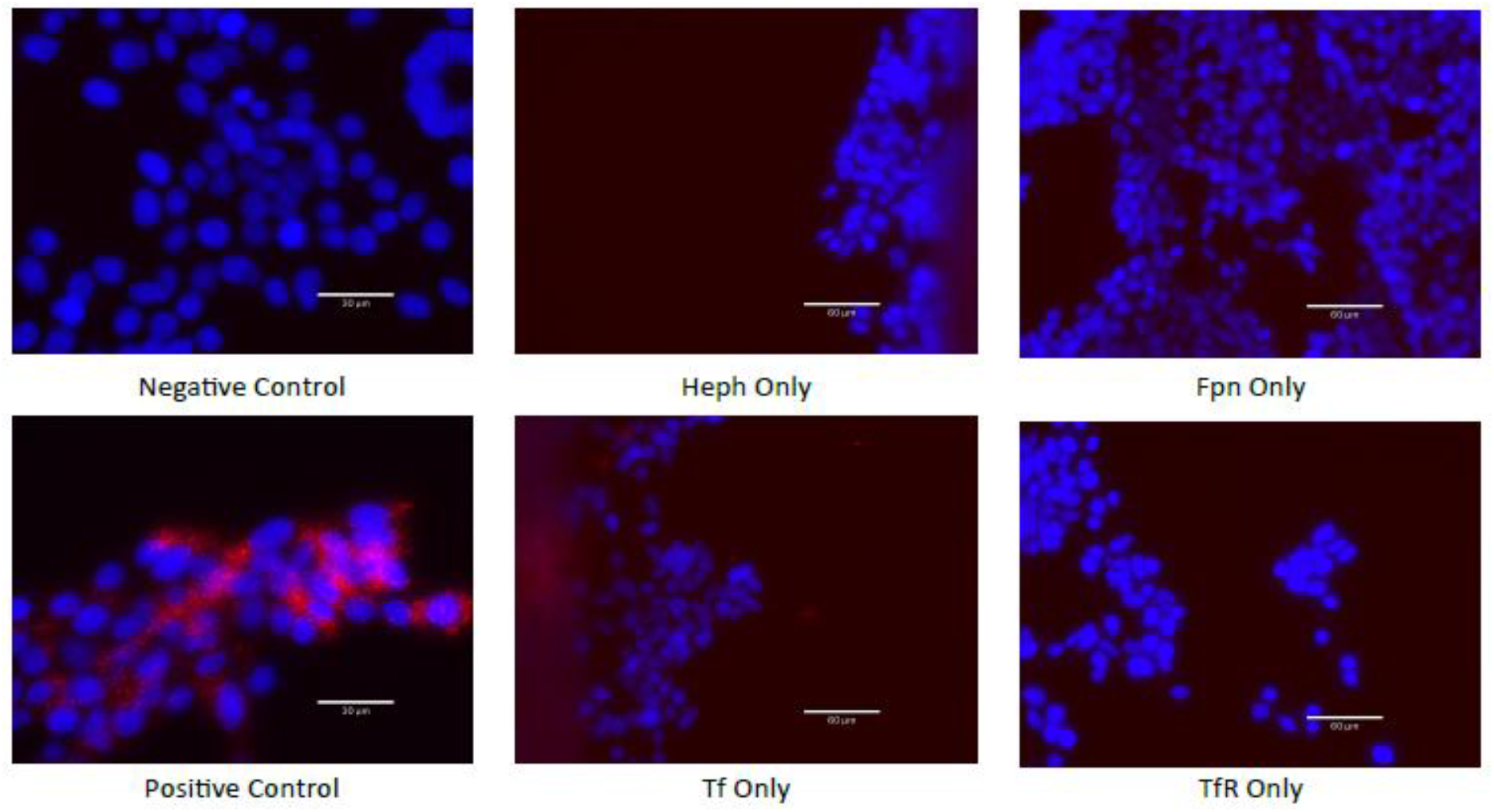
PLA Controls. A number of controls were employed while optimizing the use of PLA to determine the interactions apo- and holo-Tf had with Heph and Fpn respectively. The negative control consisted of probing for two proteins known to not interact – here MBP and ferritin. The positive control consisted of probing for two proteins known to interact – here exogenous Tf and TfR. Furthermore, each antibody used to probe for Fpn, Heph, Tf, and TfR was assessed for nonspecific binding by performing solo incubations and ensuring the antibody alone did not produce PLA signal.

**Supplemental Figure 3.**
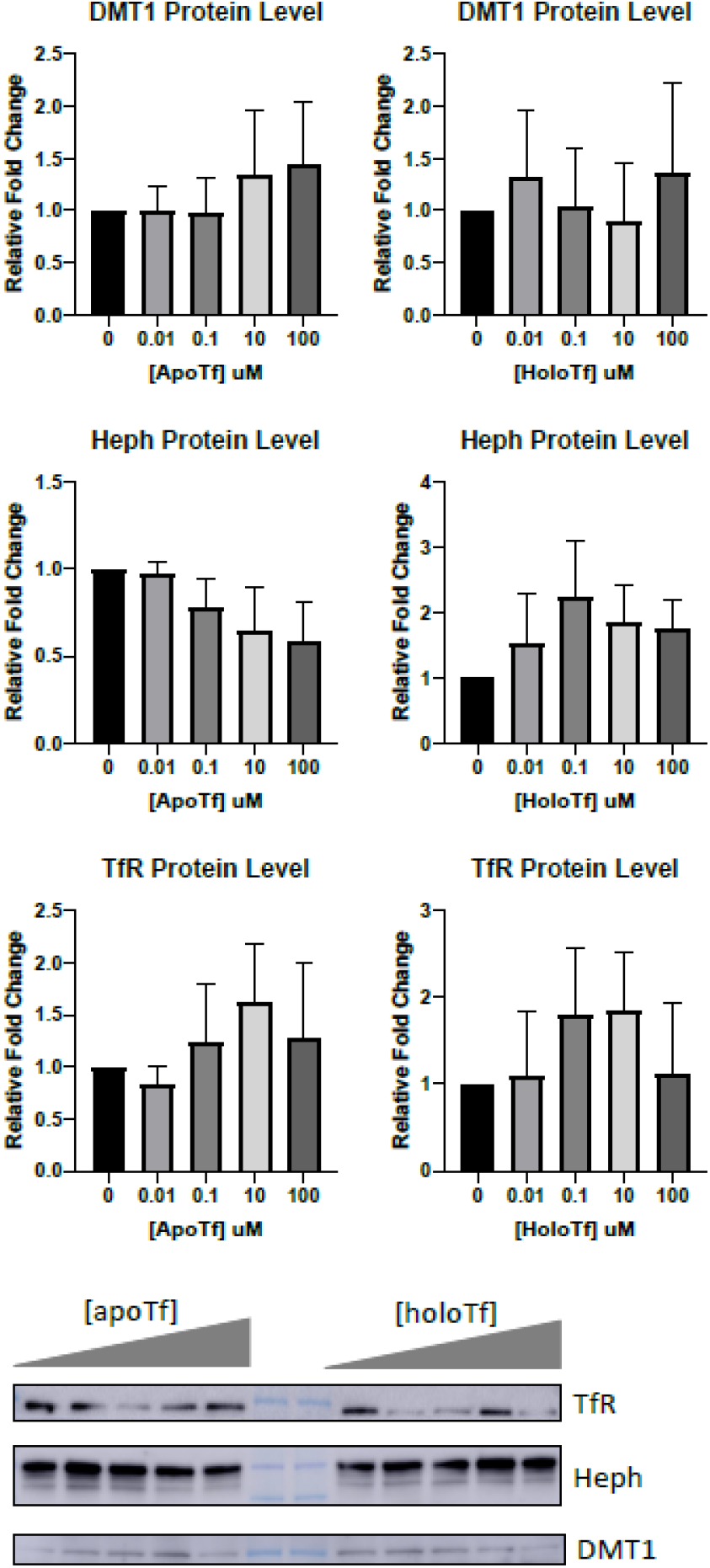
Additional Iron Regulatory Proteins with Holo-Tf Incubation. iPSC-derived ECs were incubated with holo-Tf in the basal chamber of Transwell inserts. The cells were collected and additional iron regulatory proteins were probed for using immunoblotting. Neither apo-nor holo-Tf incubations resulted in significant changes to DMT1, Heph, or TfR protein levels. n=3 for all experiments, means of biological replicates ± SEM were evaluated for statistical significance using one-way ANOVA with Tukey’s posttest for significance.

## References

1. Kim Y, Connor JR. The roles of iron and HFE genotype in neurological diseases. Mol Aspects Med 2020; 75: 100867.

2. Wade QW, Chiou B, Connor JR. Iron uptake at the blood-brain barrier is influenced by sex and genotype. Adv Pharmacol 2019; 84: 123–145.

3. Chiou B, Neal EH, Bowman AB, et al. Endothelial cells are critical regulators of iron transport in a model of the human blood-brain barrier. J Cereb Blood Flow Metab 2019; 39: 2117–2131.

4. Duck KA, Simpson IA, Connor JR. Regulatory mechanisms for iron transport across the blood-brain barrier. Biochem Biophys Res Commun 2017; 494: 70–75.

5. McCarthy RC, Kosman DJ. Mechanisms and regulation of iron trafficking across the capillary endothelial cells of the blood-brain barrier. Front Mol Neurosci 2015; 8: 31.

6. Simpson IA, Ponnuru P, Klinger ME, et al. A novel model for brain iron uptake: introducing the concept of regulation. J Cereb Blood Flow Metab 2015; 35: 48–57.

7. Baringer SL, Neely EB, Palsa K, et al. Regulation of brain iron uptake by apo- and holo-transferrin is dependent on sex and delivery protein. Fluids Barriers CNS 2022; 19: 49.

8. Beard JL, Wiesinger JA, Li N, et al. Brain iron uptake in hypotransferrinemic mice: influence of systemic iron status. J Neurosci Res 2005; 79: 254–261.

9. Connor JR, Snyder BS, Beard JL, et al. Regional distribution of iron and iron-regulatory proteins in the brain in aging and Alzheimer’s disease. Journal of Neuroscience Research 1992; 31: 327–335.

10. Dlouhy AC, Bailey DK, Steimle BL, et al. Fluorescence resonance energy transfer links membrane ferroportin, hephaestin but not ferroportin, amyloid precursor protein complex with iron efflux. J Biol Chem 2019; 294: 4202–4214.

11. Ji C, Steimle BL, Bailey DK, et al. The Ferroxidase Hephaestin But Not Amyloid Precursor Protein is Required for Ferroportin-Supported Iron Efflux in Primary Hippocampal Neurons. Cell Mol Neurobiol 2018; 38: 941–954.

12. Nemeth E, Tuttle MS, Powelson J, et al. Hepcidin regulates cellular iron efflux by binding to ferroportin and inducing its internalization. Science 2004; 306: 2090–2093.

13. Lin L, Valore EV, Nemeth E, et al. Iron transferrin regulates hepcidin synthesis in primary hepatocyte culture through hemojuvelin and BMP2/4. Blood 2007; 110: 2182–2189.

14. You L, Yu P-P, Dong T, et al. Astrocyte-derived hepcidin controls iron traffic at the blood-brain-barrier via regulating ferroportin 1 of microvascular endothelial cells. Cell Death Dis 2022; 13: 667.

15. De Domenico I, Ward DM, Langelier C, et al. The molecular mechanism of hepcidin-mediated ferroportin down-regulation. Mol Biol Cell 2007; 18: 2569–2578.

16. Eid C, Hémadi M, Ha-Duong N-T, et al. Iron uptake and transfer from ceruloplasmin to transferrin. Biochim Biophys Acta 2014; 1840: 1771–1781.

17. Sokolov AV, Voynova IV, Kostevich VA, et al. Comparison of Interaction between Ceruloplasmin and Lactoferrin/Transferrin: to Bind or Not to Bind. Biochemistry (Mosc) 2017; 82: 1073–1078.

18. Sakajiri T, Nakatsuji M, Teraoka Y, et al. Zinc mediates the interaction between ceruloplasmin and apo-transferrin for the efficient transfer of Fe(III) ions. Metallomics 2021; 13: mfab065.

19. Neal EH, Marinelli NA, Shi Y, et al. A Simplified, Fully Defined Differentiation Scheme for Producing Blood-Brain Barrier Endothelial Cells from Human iPSCs. Stem Cell Reports 2019; 12: 1380–1388.

20. Palsa K, Baringer SL, Shenoy G, et al. Extracellular vesicles are involved in iron transport from Human blood brain barrier endothelial cells and are modified by iron status. Journal of Biological Chemistry.

21. Nwogu N, Ortiz LE, Kwun HJ. Surface charge of Merkel cell polyomavirus small T antigen determines cell transformation through allosteric FBW7 WD40 domain targeting. Oncogenesis 2020; 9: 53.

22. Chiou B, Lucassen E, Sather M, et al. Semaphorin4A and H-ferritin utilize Tim-1 on human oligodendrocytes: A novel neuro-immune axis. Glia 2018; 66: 1317–1330.

23. Kondaiah P, Aslam MF, Mashurabad PC, et al. Zinc induces iron uptake and DMT1 expression in Caco-2 cells via a PI3K/IRP2 dependent mechanism. Biochem J 2019; 476: 1573–1583.

24. Qiao B, Sugianto P, Fung E, et al. Hepcidin-induced endocytosis of ferroportin is dependent on ferroportin ubiquitination. Cell Metab 2012; 15: 918–924.

25. Patton SM, Wang Q, Hulgan T, et al. Cerebrospinal fluid (CSF) biomarkers of iron status are associated with CSF viral load, antiretroviral therapy, and demographic factors in HIV-infected adults. Fluids Barriers CNS 2017; 14: 11.

26. Bergamaschi G, Villani L. Serum hepcidin: a novel diagnostic tool in disorders of iron metabolism. Haematologica 2009; 94: 1631–1633.

27. Wallace DF, McDonald CJ, Ostini L, et al. The dynamics of hepcidin-ferroportin internalization and consequences of a novel ferroportin disease mutation. Am J Hematol 2017; 92: 1052–1061.

28. Jiang R, Hua C, Wan Y, et al. Hephaestin and ceruloplasmin play distinct but interrelated roles in iron homeostasis in mouse brain. J Nutr 2015; 145: 1003–1009.

29. Yeh K-Y, Yeh M, Glass J. Interactions between ferroportin and hephaestin in rat enterocytes are reduced after iron ingestion. Gastroenterology 2011; 141: 292–299, 299.e1.

30. Ha-Duong N-T, Eid C, Hémadi M, et al. In vitro interaction between ceruloplasmin and human serum transferrin. Biochemistry 2010; 49: 10261–10263.

31. Wally J, Halbrooks PJ, Vonrhein C, et al. The crystal structure of iron-free human serum transferrin provides insight into inter-lobe communication and receptor binding. J Biol Chem 2006; 281: 24934–24944.

32. Collins JF, Wessling-Resnick M, Knutson MD. Hepcidin regulation of iron transport. J Nutr 2008; 138: 2284–2288.

33. Hänninen MM, Haapasalo J, Haapasalo H, et al. Expression of iron-related genes in human brain and brain tumors. BMC Neurosci 2009; 10: 36.

34. Vela D. Hepcidin, an emerging and important player in brain iron homeostasis. J Transl Med 2018; 16: 25.

35. Yanase K, Uemura N, Chiba Y, et al. Immunoreactivities for hepcidin, ferroportin, and hephaestin in astrocytes and choroid plexus epithelium of human brains. Neuropathology 2020; 40: 75–83.

36. Raha-Chowdhury R, Raha AA, Forostyak S, et al. Expression and cellular localization of hepcidin mRNA and protein in normal rat brain. BMC Neurosci 2015; 16: 24.

37. Zechel S, Huber-Wittmer K, von Bohlen und Halbach O. Distribution of the iron-regulating protein hepcidin in the murine central nervous system. J Neurosci Res 2006; 84: 790–800.

38. Xu Y, Zhang Y, Zhang J-H, et al. Astrocyte hepcidin ameliorates neuronal loss through attenuating brain iron deposition and oxidative stress in APP/PS1 mice. Free Radical Biology and Medicine 2020; 158: 84–95.

39. McCarthy RC, Kosman DJ. Glial cell ceruloplasmin and hepcidin differentially regulate iron efflux from brain microvascular endothelial cells. PLoS One 2014; 9: e89003.

40. Enculescu M, Metzendorf C, Sparla R, et al. Modelling Systemic Iron Regulation during Dietary Iron Overload and Acute Inflammation: Role of Hepcidin-Independent Mechanisms. PLoS Comput Biol 2017; 13: e1005322.

41. Lane DJR, Ayton S, Bush AI. Iron and Alzheimer’s Disease: An Update on Emerging Mechanisms. Journal of Alzheimer’s Disease 2018; 64: S379–S395.

42. Du G, Wang E, Sica C, et al. Dynamics of Nigral Iron Accumulation in Parkinson’s Disease: From Diagnosis to Late Stage. Mov Disord. Epub ahead of print 25 May 2022. DOI: 10.1002/mds.29062.

43. Ji C, Kosman DJ. Molecular mechanisms of non-transferrin-bound and transferring-bound iron uptake in primary hippocampal neurons. J Neurochem 2015; 133: 668–683.

44. Erikson KM, Pinero DJ, Connor JR, et al. Regional Brain Iron, Ferritin and Transferrin Concentrations during Iron Deficiency and Iron Repletion in Developing Rats. The Journal of Nutrition 1997; 127: 2030–2038.

45. Beard JL, Wiesinger JA, Li N, et al. Brain iron uptake in hypotransferrinemic mice: influence of systemic iron status. J Neurosci Res 2005; 79: 254–261.

46. Felt BT, Lozoff B. Brain iron and behavior of rats are not normalized by treatment of iron deficiency anemia during early development. J Nutr 1996; 126: 693–701.

